# Composition and higher-order structure in nucleic acids sequenced from a chondrite

**DOI:** 10.64898/2026.01.26.701670

**Authors:** Carmel Farage, George M. Church, Ido Bachelet

## Abstract

The known tree of life occupies an infinitesimal region of the space of all mathematically possible evolutionary histories, yet our sequence analysis frameworks are implicitly calibrated to it and to its associated compositional and grammatical regularities. Here we analyze nucleic acid molecules sequenced from the Zag meteorite as part of a broader effort to understand how nucleic acid sequence composition and higher-order structure are shaped under chemically divergent environments. We characterize these sequences across multiple analytical layers, and show that they lack signatures of protein-coding organization, translational periodicity, or known biological grammar. At the same time, they deviate significantly from random or composition-only null models, displaying constrained complexity and low-dimensional structure in *k*-mer frequency space. Multiple tests place amplification and sequencing-driven artifacts and metagenomic contaminants at a low likelihood. Taken together, these findings indicate that the Zag sequences occupy an unusual region of sequence space that is not readily accounted for by known biological or technical models, thereby narrowing, but not resolving, the range of plausible explanations and motivating independent replication and further investigation.

## Introduction

It is generally assumed that the currently known tree of life (ToL) accounts for only a small fraction of all living organisms on earth^1^. Estimates of the actual terrestrial biodiversity vary widely, but even under conservative assumptions, e.g. that the known fraction represents 0.1-1.0% of extant life, the complete ToL would still occupy a negligible fraction of the space of possible DNA sequences with a comparable length distribution. For example, the space of all DNA sequences of length ∼ 100 bases contains 4^100^ (∼ 10^60^) distinct sequences, whereas even a very conservative estimate of the total number of nucleotides represented in a complete ToL is on the order of 10^17^-10^18^. While biological function and evolutionary history strongly constrain which regions of sequence space are accessible, this disparity in scale implies that, in principle, the sequence space could contain an enormous number of independent evolutionary histories - trees of life occupying largely disjoint regions of sequence space and sharing no common ancestry.

New sequences are usually compared to the known database in search for novel genetic material using tools such as BLAST^2^ and Kraken2^3^, with mapping below a certain threshold indicating that the material is indeed novel. However, it is crucial to recognize that the validity of this practice is predicated upon the assumption that any newly discovered sequence must originate from the ToL i.e. from some organism within it, known or yet unknown. This assumption is arguably scientifically rigorous, but it might also bias our conclusions in the face of hypothetical DNA or RNA originating from a system that is foreign to the one we know.

Much of the newly found genetic material is regarded as novel according to current methods for measuring novelty. This state of matters is suitable for the taxonomic project of filling the blanks in the ToL. However, in terms of detecting foreignness, it has little value as even very novel genetic material would be considered a yet-unknown terrestrial organism or a variant of existing content. Even the most distant branches of the ToL appear to share more similarity than difference^4–9^.

We recently completed a collaborative effort that resulted in a database of nucleic acid molecules sequenced from internal samples from two masses of the Zag chondrite^10^, a fragment of a H-type ordinary chondrite that fell on earth in 1998 and was reported to contain several compositional peculiarities. A stringent, multi-method filtering of sequences that mapped to known material highlighted a fraction (totaling approximately 17,000 reads) of unmapped material, which was the focus of this work.

## Results

The meteorite known as Zag is a fragment of a H3-6 chondrite, from a yet-unclear parent body^11,12^, that fell in Western Sahara near Zag, Morocco, in August 1998^10^. Analyses of the meteorite revealed that it contains inclusions of liquid water within microscopic halite crystals^13^, and that it formed approximately 4.7 Billion years ago based on ^129^I-^129^Xe radiometric dating^14,15^. More recently, complex organic molecules have been reported from inside the chondrite^16–18^, which contains micrometer-sized organic matter aggregates that are rare in other H-type meteorites^17^. C-type meteorites such as Murchison, Murray, and Orgueil, have long been known to contain basic building blocks of life, including amino acids^19,20^ and nucleobases^21–23^. We were interested in the possibility of identifying more complex nucleic acids in two Zag masses, weighing 18.8 and 1.48 kg **(Fig. 1A, 1B)**.

**Figure 1.**
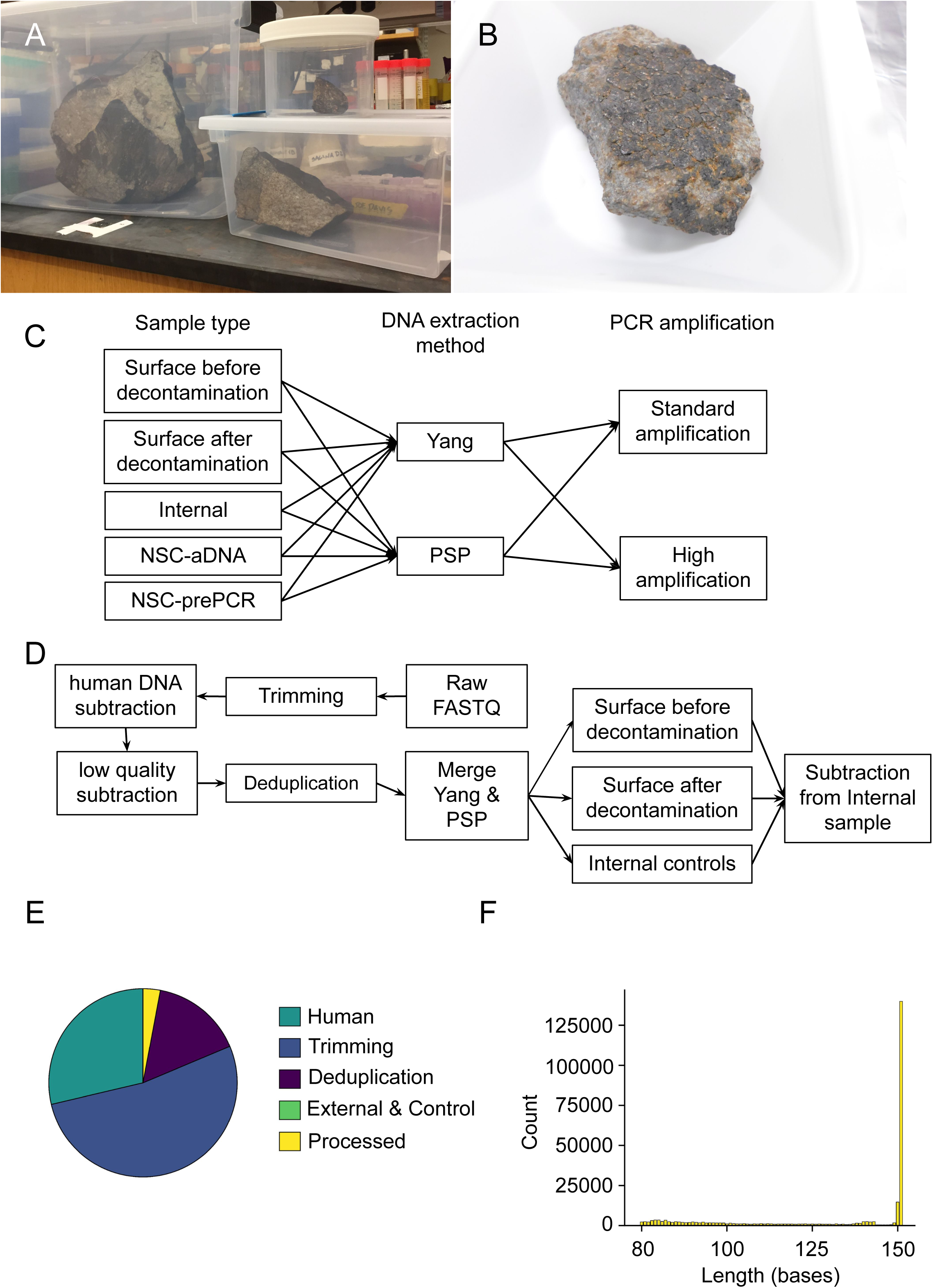
(A, B) The two Zag meteorite masses (18.8 kg and 1.48 kg). (C–F) Sample preparation and sequencing pipeline. Three sample types were collected: surface (before decontamination), surface after decontamination, and internal fracture surface. DNA was extracted using Yang-urea and PowerSoil Pro methods. Two blunt-end libraries per extract were prepared with standard and high amplification levels. All data underwent multi-stage QC: adapter trimming, quality filtering, removal of human reads (∼89% and ∼86% removed for R1/R2), de novo deduplication (∼5% R1, ∼4% R2 retained), and subtraction of reads detected in surface samples, no-sample controls, and pre-PCR controls. Internal read sets were mapped to ChocoPhlAn and BLAST NT; unmapped reads (9,252 high-amp and 8,083 standard-amp) formed the final dataset. Scale bars and sample locations as indicated.

The sample preparation and sequencing workflow is depicted in **Fig. 1C-1F**. Three samples were collected: (1) from the meteorite surface before decontamination (Surface), (2) from the surface after decontamination (Surface-decontaminated) which removed DNA present on the surface, and (3) from a fresh internal surface of the meteorite (Internal) after fracturing the specimen in a clean room. DNA was extracted from each sample using two different methods, Yang-urea ancient DNA extraction method (Yang), and QIAGEN DNeasy PowerSoil Pro kit (PSP). Two blunt-end DNA libraries were prepared from each extract. The two libraries were prepared using different levels of DNA amplification – standard and high. All sequence data were subjected to a multi-stage quality control and subtraction pipeline prior to downstream analysis. Adapter trimming, quality filtering, and removal of human reads resulted in the exclusion of the majority of raw reads, with approximately 89% and 86% of the R1 and R2 pair reads removed at this stage, consistent with substantial background contamination expected in meteoritic samples handled in terrestrial environments. Following de novo deduplication, approximately 5.1% (R1) and 4.2% (R2) of read pairs remained. To operationally define sequences associated with the internal fracture surface of the Zag chondrite, we further subtracted all reads detected in surface samples (pre- and post-decontamination), no-sample controls, and pre-PCR controls. This subtraction was performed independently for standard- and high-amplification libraries, yielding a conservative “internal” read set defined by absence from all control contexts rather than by any positive taxonomic assignment. Reads in the internal sets were first mapped to curated genome collections using ChocoPhlAn, followed by alignment against the BLAST NT database. A substantial fraction of reads mapped broadly to known bacterial clades, indicating that the pipeline successfully detects conventional terrestrial signals when present. However, a small subset of reads remained unmapped after both steps, comprising 9,252 reads in the high-amplification library and 8,083 reads in the standard-amplification library. These unmapped reads (**Supplementary files 1-2)** were retained as a pooled set for further analysis without assuming novelty of origin, but rather as sequences not readily assignable within current reference frameworks under stringent alignment criteria.

Base composition analysis showed a moderate AT enrichment (A 28%, T 29%, G 23%, C 20%), with no significant C-deamination^24^. Given an activation energy (for C-deamination in single-stranded DNA at 37 °C) of 117 KJ/mol and a rate constant of 10^-10^ s^-1^, by Arrhenius extrapolation, C-deamination in space (2.7 °K) has a calculated rate constant on the order of ∼ 10^-2030^, equivalent to zero deamination over geological time scales, in accordance with the observed base composition.

We next evaluated whether the sequences could represent fragments of conventional protein-coding genomes. For this we curated 5 control groups (all n = 17,000): raw Illumina sequencing files of *E. coli* coding and non-coding/intergenic sequences; a group of reads rejected from a standard *E. coli* Illumina sequencing run; a group of reads from the pre-filtered Zag sequences; and a group of reads from the Zag dataset that was filtered out. Triplet periodicity analysis in all 6 frames showed no detectable signal **(Fig. 2A)**. Frame-aware hexamer log-odds scoring robustly separated coding from noncoding controls, with the Zag sequences aligning with noncoding and rejected datasets rather than coding sequences, indicating a lack of codon-pair structure characteristic of translation **(Fig. 2B)**. Stop-codon avoidance analysis revealed that the sequences exhibited stop frequencies indistinguishable from dinucleotide-preserving shuffles **(Fig. 2C)**. Scans across a range of periods (p = 2 to p = 8) failed to reveal periodic signals **(Fig. 2D, 2E)**, as did a more sensitive analysis specifically for quadruplet periodic structures **(Fig. 2F)**.

**Figure 2.**
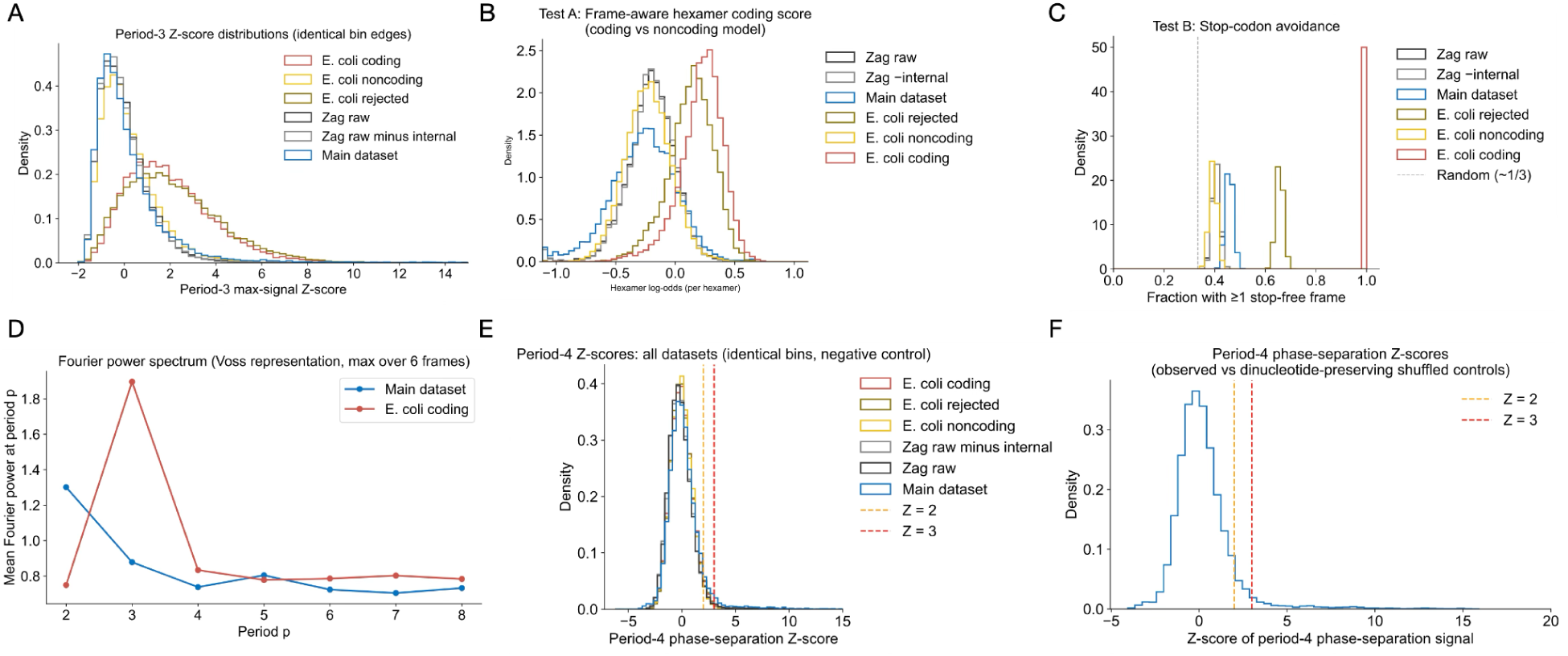
Lack of protein-coding and periodic structure in Zag sequences. Six control datasets (n=17,000 each, main dataset n=17,335): E. coli coding, E. coli noncoding, E. coli rejected, Zag raw, Zag raw minus internal, and main (internal) Zag dataset. (A) Triplet periodicity in all six frames: no detectable signal in Zag; period-3 Z-scores are not elevated relative to null. (B) Frame-aware hexamer log-odds score: main dataset mean = −0.505, median = −0.438, fraction score > 0 = 6.1%; E. coli coding mean = 0.455, fraction > 0 = 96.9%. Zag aligns with noncoding and rejected controls, not coding. ROC AUC (hexamer, coding vs noncoding) = 0.964. (C) Stop-codon avoidance: Zag stop frequencies indistinguishable from dinucleotide-preserving shuffles. (D, E) Periodicity scans (p=2–8): no periodic signals. (F) Period-4 (quadruplet) phase-separation Z-scores (negative control, 200 dinucleotide-preserving shuffles per sequence): main dataset mean Z = 0.225, median Z = −0.021, fraction Z ≥ 2 = 7.7%, fraction Z ≥ 3 = 3.9%; values are not elevated relative to null (ecoli_coding mean Z = −0.015; ecoli_noncoding mean Z = −0.043). No period-4 signal. Together, these results exclude protein-coding organization and translational or other periodic structure.

We asked whether the sequences behave as random or weakly constrained polymers. Analysis of 3-mer Shannon entropy revealed values significantly lower than those obtained from mono- and dinucleotide-preserving shuffled controls, indicating a non-random sequence structure **(Fig. 3A, 3B, 3C)**. Enrichment of simple homopolymers and short tandem fragments was observed **(Fig. 3D, 3E)**, with no systematic enrichment of palindromic or hairpin-forming sequences **(Fig. 3F)**.

**Figure 3.**
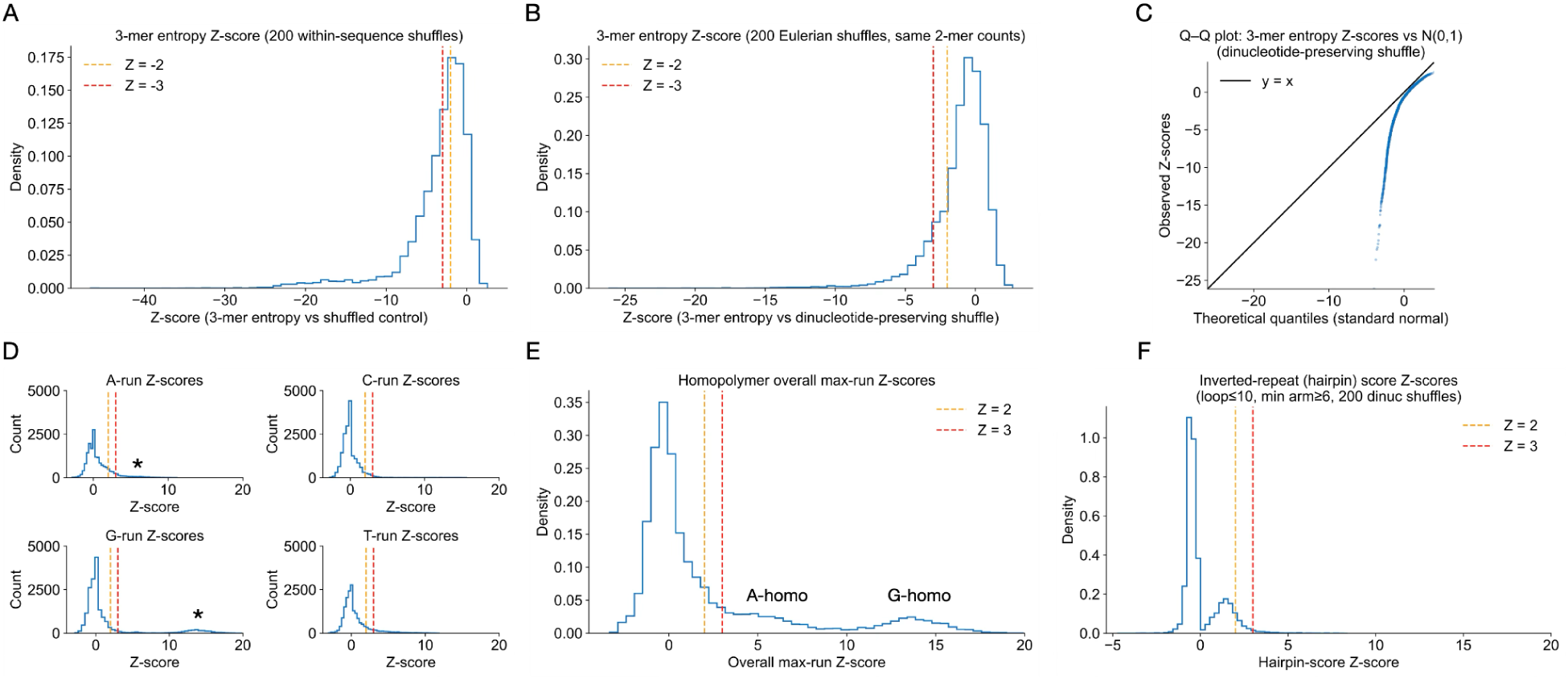
Non-random sequence complexity and motif structure. (A–C) 3-mer Shannon entropy versus mono- and dinucleotide-preserving shuffled controls (200 shuffles per sequence). Main dataset mean 3-mer entropy = 4.96; E. coli coding = 5.40; E. coli noncoding = 5.35. Zag sequences exhibit significantly lower entropy (Z-scores strongly negative vs shuffled null), indicating constrained, non-random structure. (D, E) Global 3-mer and 4-mer enrichment/depletion (Z-scores vs dinucleotide-preserving shuffle, n=200): strong enrichment of homopolymers and tandem motifs—e.g. GGGG fold-change 1.80, Z = 198; GGG fold-change 1.22, Z = 123; GTGT fold-change 1.42, Z = 62; ACAC fold-change 1.50, Z = 58. Top depleted 3-mers include GGT (fold 0.75, Z = −66), CAA (fold 0.82, Z = −59). Asterisks in D denote A-homo and G-homo peaks at E. (F) Palindrome and hairpin-forming sequences: no systematic enrichment. The combination of reduced entropy and homopolymer/tandem enrichment is consistent with constrained, non-enzymatic generation rather than biological genomes or pure random polymers.

To evaluate likelihood for the existence of grammatical structures, first- and second-order Markov models were used to compute log-likelihood ratio (LLR) scores for all reads, quantifying how strongly each sequence conforms to biological coding grammar. Both models yielded consistent results, with the main dataset exhibiting a unimodal LLR distribution that is clearly shifted away from coding-like sequences and shows no evidence for a distinct PCR-artifact–specific subpopulation **(Fig. 4A, 4B)**. In parallel, we computed a hexamer-based coding score for each sequence **(Fig. 4C**; ROC curves for Markov & hexamer models are described in **Fig. 4D)**. As expected, biological sequences collapse onto a narrow, low-dimensional manifold in joint hexamer–Markov space, whereas the Zag dataset exhibits multiple manifolds, consistent with heterogeneous sequence generation processes, constraints, and rules that do not conform to a single grammatical model **(Fig. 4E)**.

**Figure 4.**
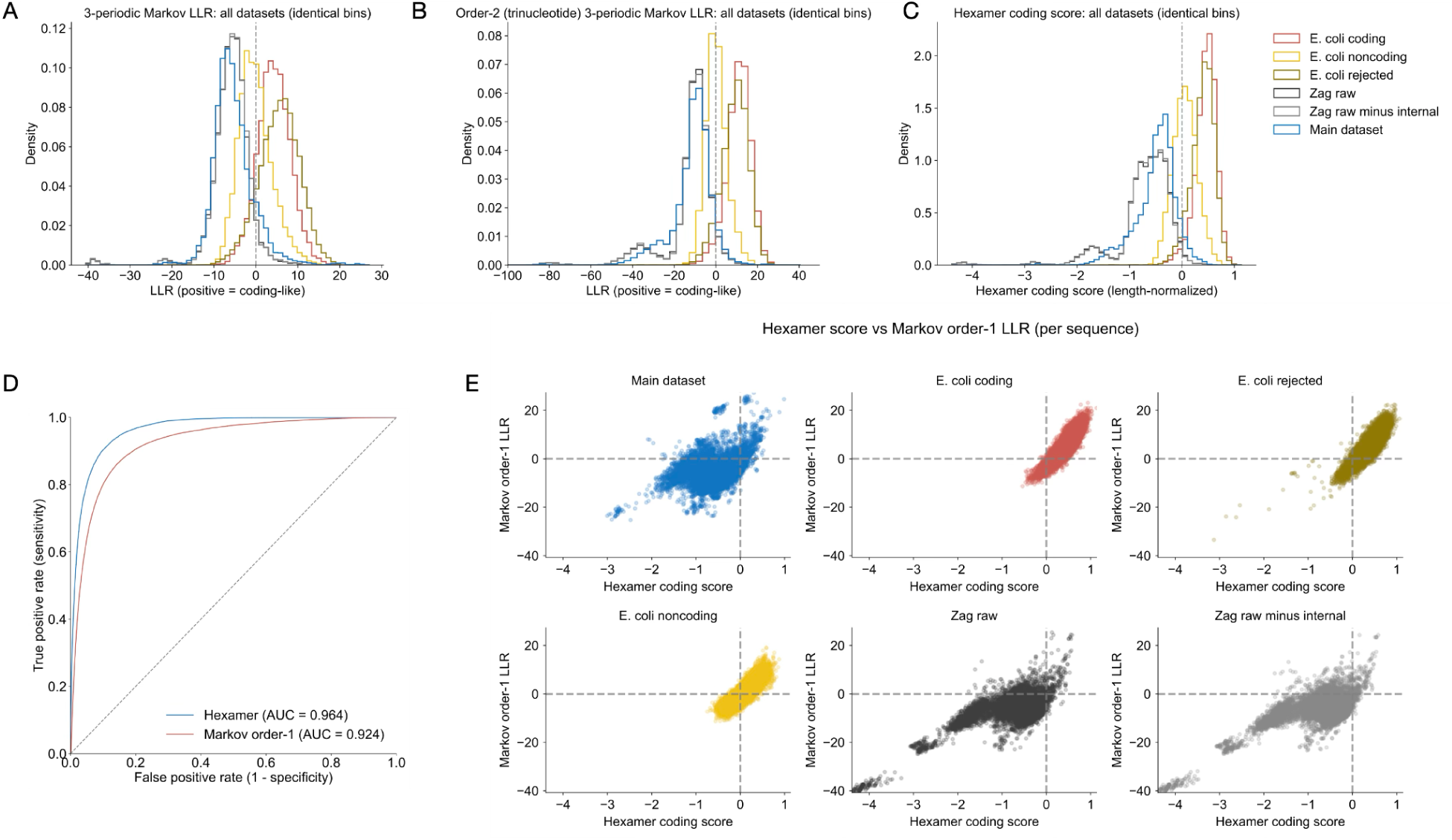
Grammar space: Markov and hexamer models. (A, B) First- and second-order Markov log-likelihood ratio (LLR) distributions. Main dataset: order-1 mean LLR = −5.27, median = −5.96, fraction LLR > 0 = 12.2%, fraction LLR > t95 (95% specificity) = 3.7%; order-2 mean LLR = −11.46, median = −9.79, fraction LLR > 0 = 6.8%. E. coli coding: order-1 mean LLR = 4.37, fraction > 0 = 87.5%; order-2 mean LLR = 10.95. ROC AUC (order-1, coding vs noncoding) = 0.917; ROC AUC (order-2) = 0.956. Zag exhibits unimodal, negative LLR distributions, shifted away from coding-like sequences with no distinct PCR-artifact subpopulation. (C) Per-sequence hexamer coding score. (D) ROC curves for Markov and hexamer models. (E) Joint hexamer–Markov space: biological controls collapse onto a narrow manifold; Zag displays multiple manifolds consistent with heterogeneous, non-grammatical sequence generation under partial, orthogonal constraints rather than a single coding grammar.

We then examined the population-level structure of the dataset to determine whether it reflects a mixture of multiple discrete sources, such as a metagenomic assemblage, or a single homogeneous population. Gaussian mixture modeling (GMM) favored a unimodal distribution under Bayesian information criterion **(Fig. 5A)**, a result independently supported by Hartigan’s dip test on the primary principal component **(Fig. 5B)**. While the dataset displays heterogeneity in higher-order grammar space, unimodality along the dominant principal axis indicates that the sequences are best explained by a single underlying generative process operating under multiple constraints, rather than by a mixture of unrelated biological or technical sources.

**Figure 5.**
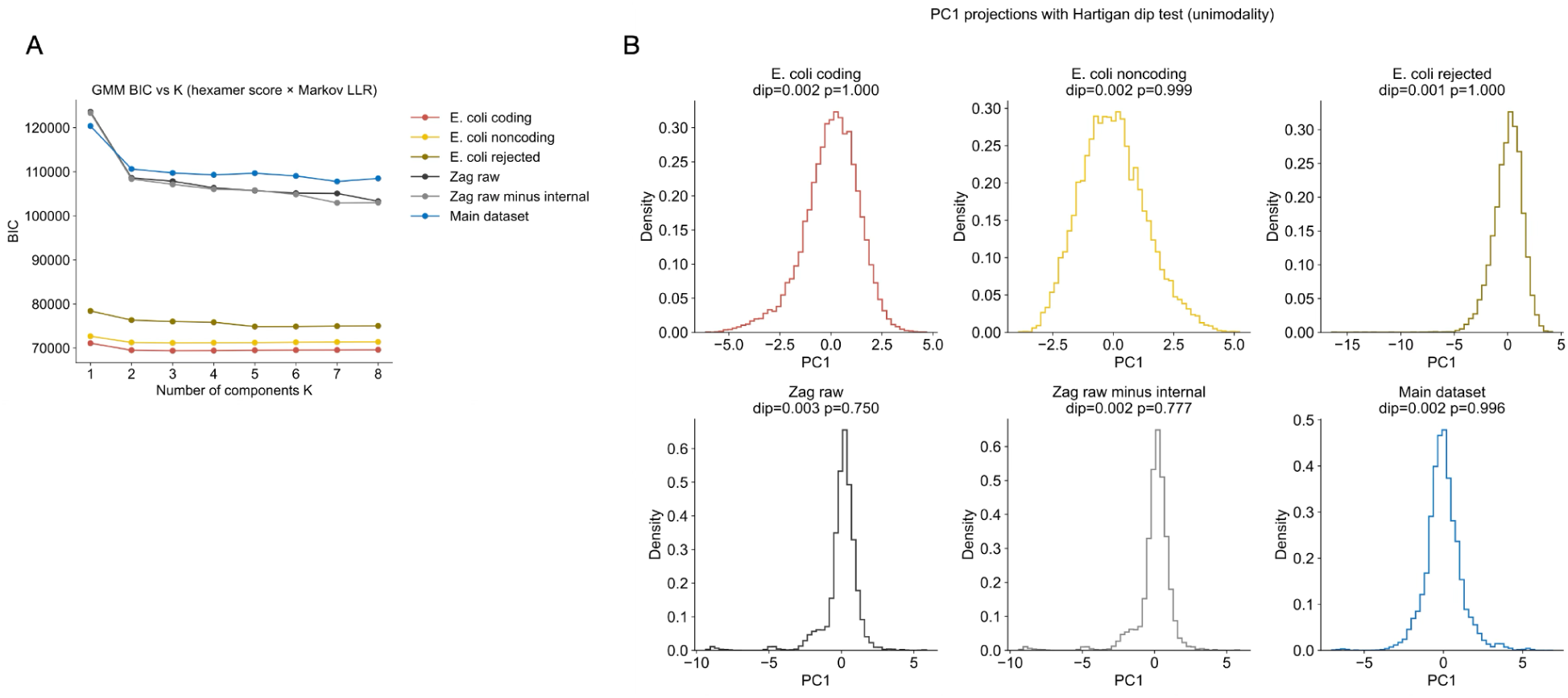
Population structure: unimodality along dominant variance. (A) Gaussian mixture model (GMM) BIC versus number of components (K=1–8) in hexamer–Markov space. Main dataset: selected K* = 7, BIC_min = 107,789, GMM weight entropy = 1.60. E. coli coding K* = 3; ecoli_rejected K* = 5; zag_raw K* = 8. (B) Hartigan’s dip test on PC1 (2000 bootstrap samples, seed 42). Main dataset: dip statistic = 0.0018, p = 0.996; ecoli_coding p = 0.9995; ecoli_rejected p = 1.000; ecoli_noncoding p = 0.9985; zag_raw p = 0.75; zag_raw_minus_internal p = 0.78. High p-values support unimodality. Together, these indicate a single underlying generative process for the main dataset rather than a mixture of discrete sources (e.g. metagenomic assemblage), despite heterogeneity in grammar space.

Given the low-input nature of the material, we tested whether amplification and technical sequencing artifacts could account for the observed sequences. We first assessed library-end or adapter-related effects, using a reference set of Illumina and Nextera library oligonucleotides. 13.5% of sequences showed reliable match to bona fide technical oligonucleotides. In order to obtain an estimated upper bound to such contaminations, we ran a deliberately permissive screen based on exact or near-exact short substring matches (minimum match length 10 nt), with ∼ 32.66% showing some similarity to known adapter or primer sequences **(Fig. S1)**. These searches show that even under extremely stringent conditions, roughly ⅔ of the dataset cannot be reliably explained as known technical artifacts, with the real number likely closer to ∼ 90%. The estimated chimera rate was below 0.1%. Edit-distance analyses revealed no amplification-driven “error halo,” with most reads having no close variants and no dominance of parent-derived families. Rank–abundance distributions were dominated by singletons rather than exhibiting power-law or geometric decay, and amplification jackpot metrics were minimal, with a Gini coefficient of 0.014. Collectively, these metrics place PCR-driven artifact generation at a low likelihood as the source of the dataset. We further performed principal component analysis of 6-mer frequency profiles, revealing a low-dimensional manifold structure rather than a diffuse cloud expected from random noise or amplification artifacts. Consistently, MinHash-based Jaccard similarity between the dataset and dinucleotide-preserving shuffled controls was on the order of 10^-3^, several orders of magnitude lower than null expectations. Side-by-side comparisons with explicitly constructed null datasets demonstrated that random noise, PCR artifacts, and composition-only models fail to reproduce the observed structure.

In order to analyze the higher-order *k-*mer structure in this material, we first characterized the basal layer of *k*-mers in the ToL using the BLAST NT database as a source. This database is a non-redundant collection of diverse nucleotide sequences from various sources, including GenBank, EMBL, DDBJ, PDB, and RefSeq^25^. The NT database version used in this study was downloaded in December 2023 and at that point consisted of 102,960,590 records. We calculated *k*-mer representation for *k* = 12, 14, 16, 18, 20, and 21 in decreasing NT subsample sizes and observed a pattern which is in agreement with previous findings^26^ **(Fig. 6A)**. At *k* = 12, 14, and 16, 100% of the *k*-mers were represented in NT; at *k* = 20, only 29.24% of all possible *k*-mers (4^20^ = 1.099×10^12^) were found to be represented in NT; and at *k* = 21, only 10.04% of all possible *k*-mers (4^21^ = 4.398×10^12^) were represented in NT. We defined a parameter, RD_50_, which equals the database size at which 50% of the *k*-mers (for a certain *k*) are represented, and observed a significant correlation between RD_50_ and *k* **(Fig. 6B)**, suggesting that all 20-mers and 21-mers might be represented once NT reaches a size of ∼ 3.62×10^16^ and ∼ 1.37×10^17^ bases, respectively. The representation of 20-mers across the ToL exhibited variation between clades, in a manner suggesting that taxonomic association to a specific clade could be possible for newly found material based on its *k*-mer representation **(Fig. 6C)**. In addition, we generated various reference groups and measured their representations in NT: (1) a sample of 1,000 sequences from NT, which by definition exhibits 100% representation; (2) a sample of 1,000 randomly-generated sequences, showing average representation of 29.37%, which is approximately equal to the total fraction of 20-mers represented in NT (29.24%); (3) a previously-reported bacterial genome from a potentially new phylum^27^, representing novel genetic material, showing average representation of 52.84%. (4) an artificial “foreign” sequence composed solely from unrepresented 20-mers such that every newly created 20-mer is also unrepresented; this sequence thus exhibited 0% representation and did not map to NT. (5) the foreign sequence was reshuffled to create a fifth reference group, which showed 20% representation. The remaining set of sequences contained >1,000,000 overlapping *k-*mers (*k =* 21), of which approximately 33% were unique. Analysis of these *k*-mers revealed 100% of them to be primes, i.e. 0% of these *k*-mers were represented in the NT database, which was different than the reshuffled 20-mer prime-constructed group (20%) and the random representations of 21-mers (10.04%). **Figure 6D** places this result within a range of estimated representations^28–30^ and the empirical ones described above.

**Figure 6.**
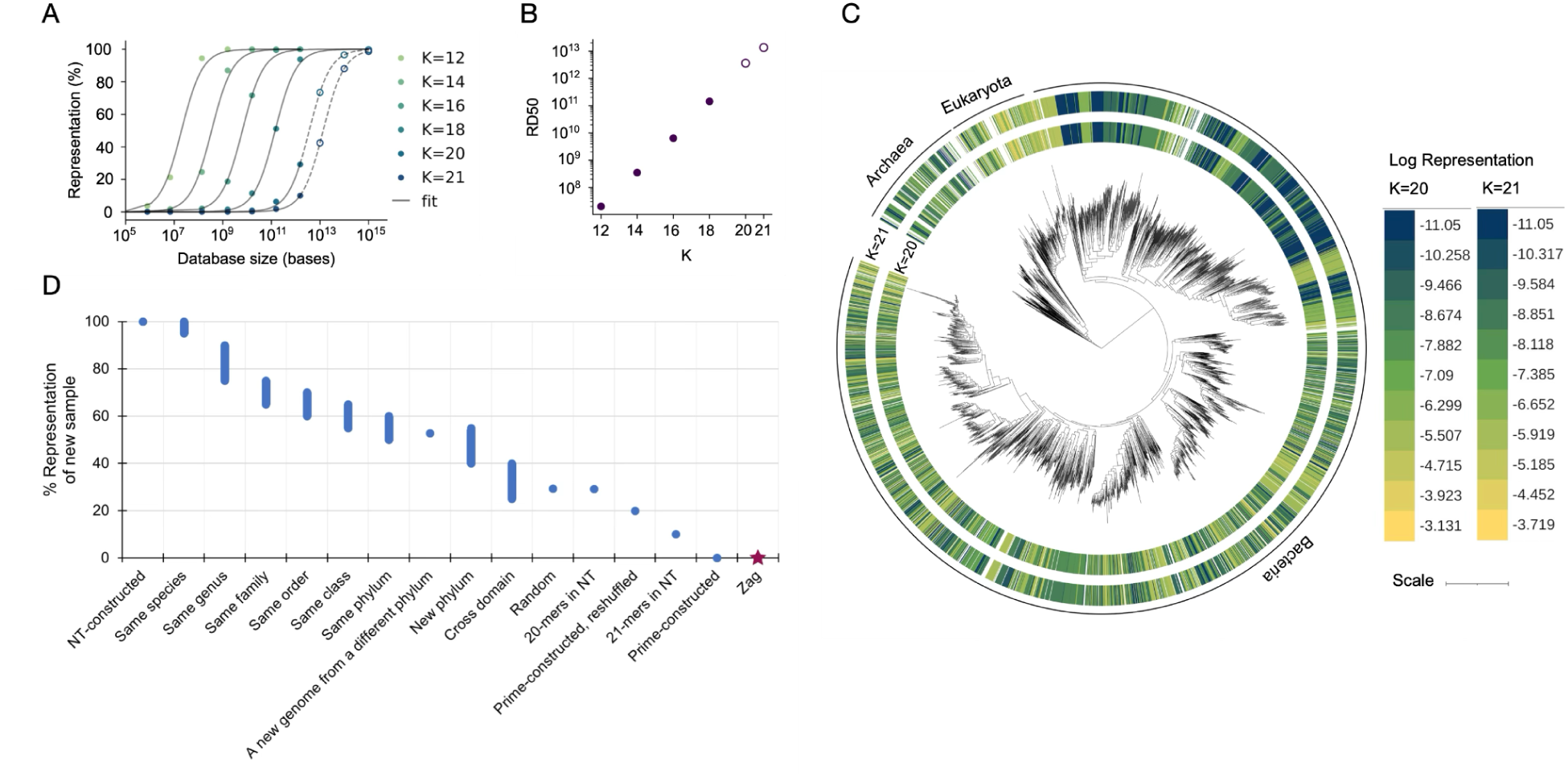
*k*-mer representation in the nucleotide database and placement of Zag sequences. (A) Fraction of possible *k*-mers represented in BLAST NT at *k*=12–21 across subsample sizes. At *k*=12–16, 100% represented; at *k*=20, 29.24% (4^20 ≈ 1.10×10^12 possible); at *k*=21, 10.04% (4^21 ≈ 4.40×10^12). (B) RD50 (database size in bases at which 50% of *k*-mers are represented) versus k; strong correlation. (C) 20-mer representation across tree-of-life clades. (D) The representation of various samples in NT, based on their *k-*mer composition. Zag 21-mers: 100% primes (0% represented in NT), distinct from reshuffled prime-built sequences (20%), random (10.04%), and novel bacterial material (∼53%). Zag occupies an extreme region of *k*-mer space consistent with absence from known biological sequence databases.

**Table S1** describes the list of analytical tests and their summarized findings.

## Discussion

This work describes a dataset of nucleic acid molecules recovered from an internal sample of the Zag chondrite and analyzed using a broad set of complementary sequence-level tests. We examined higher-order *k*-mer organization, potential coding structure, translational periodicity, grammatical coherence, sequence complexity, population structure, and susceptibility to amplification or sequencing artifacts. Across these largely independent dimensions, the sequences consistently lack hallmarks of biological genomes, including open reading frame structure, codon periodicity, and a unified coding grammar, while at the same time deviating from random or composition-only null models. While analyses of coding potential directly exclude protein-coding genomic contamination, additional population-level, grammatical, and *k*-mer based tests argue against a dominant contribution from genomic non-coding regions. Population-level analyses further distinguish the dataset from a conventional metagenomic mixture: unimodality along dominant variance axes excludes discrete genome-scale sources, whereas joint grammar-space analyses reveal heterogeneous subpopulations shaped by partial, orthogonal constraints rather than a single generative process.

Extensive controls placed technical and PCR-driven artifacts at low likelihood. In our attempt to match the dataset with sequencing and amplification artifacts, we noticed that a significant fraction of the reads exhibited extended homopolymer or low-complexity/tandem motifs. Long homopolymer tracts and strong motif biases are rare in terrestrial biological DNA and are therefore routinely treated as artifacts in conventional genomics pipelines; however, such assumptions are specific to enzymatic, template-guided replication and need not apply to non-enzymatic or abiotic polymerization processes. Under chemical models of template-independent nucleotide assembly, such as surface-catalyzed growth, base-stacking driven extension, or monomer availability constraints, low sequence complexity and homopolymer enrichment are expected rather than anomalous. Consistent with this, the sequences exhibit reproducible, non-random structure across multiple independent grammatical and statistical tests while failing all known biological coding signatures. Taken together, these observations argue in favor of non-biological DNA generated by constrained, non-enzymatic processes.

This work elaborates on an initial survey of *k-*mer structure in the material as a first analytical filter. *k*-mers have been used as the basis for various bioinformatic analysis methods and tools as they enable improvement in computational efficiency and speed^31^. We have been previously applying *k*-mer analysis to DNA and RNA obtained from chemically divergent and biologically extreme environments. In this context, the choice of *k* is of significant importance as it defines the thresholds for proximity and foreignness. As mentioned above, all *k*-mers up to 16 are already known in the database. 17 is the first value of *k* to date at which some *k*-mers are primes. We used *k* of 20 and 21 for specific reasons. 20 is the value that created a sufficiently large pool of primes from which to build “foreign” reference sequences. On the other hand, evidence suggests that *k* = 21 is optimal for comparison across genomes as it balances *k*-mer sharing and specificity^26,32,33^, however 20 and 21 are considered similar in terms of sensitivity and precision^32^.

A value for *k* could be hinted at from a different direction: what is the minimal length of biological meaning? Short, non-coding RNAs. e.g. miRNAs, which are around 20-22 nt in length, are usually considered to mark the short end of the spectrum^34,35^. Shorter RNA fragments have also been reported^36–39^, but the shortest ones (below 17 nt) may still correspond to degradation products. Very short RNAs are known to be functional, for example the 5-nt minuscule ribozyme^40,41^, but these are arguably chemical entities that are not encoded as biological messages. Altogether it is interesting that, according to present understanding, 20- and 21-mers may represent (or be proximal to) a boundary between biological and chemical meaning, which is intricately linked to the question of the potential origin of found material.

As suggested above, hypothetical origins of nucleic acids outside the ToL should not be ruled out completely, with one such source being abiotic synthesis. Although the chemistry of complete abiotic synthesis of nucleic acids is not well understood and has not been fully recreated to date despite significant progress^42–45^, it is hypothesized to have occurred on the primitive earth, although the nature of the first nucleic acid species is also unclear^46,47^. Simple precursor molecules for chemical nucleic acid synthesis are abundant in the universe, and the chemistry for their generation has been described and extensively studied in the past 70 years. The synthesis of ribose, one of the major gaps in our understanding of abiotic nucleic acid synthesis, has been suggested to occur from formaldehyde in the presence of borate and silicate minerals^48–51^; and, while boron in relatively rare in the universe (abundance is approximately 6 orders of magnitude lower than that of carbon and oxygen, and 10 orders of magnitude lower than that of hydrogen), it has been identified in local concentrates in space^52^. Chondrites typically contain boron in the range of 0.1-1 ppm^53,54^. It is not suggested here that nucleic acids are synthesized de-novo inside chondrites: this is unlikely^55^. Rather, we reiterate the hypothesis that there are yet-unknown chemical conditions under which the abiotic synthesis of nucleic acids - regardless of their base and backbone chemistry, chirality, etc. - can proceed forward from simple, abundant precursors, on earth and elsewhere in the present and past. In this context, it is intriguing to hypothesize that, since Taq polymerases exhibit inherent, low-level RNA-dependent DNA polymerase activity^56^, the Zag source material was in fact RNA, which has a clear abiotic synthesis route.

Taken together, the findings described in this paper indicate that the Zag sequences occupy an unusual region of sequence space that is not readily accounted for by known biological or technical models, thereby narrowing, but not resolving, the range of plausible explanations and motivating independent replication and further investigation. The findings also highlight the possibility that our sequence analysis tools may be inadequate to identify the products of abiotic synthesis of nucleic acids, should such hypothetical products be captured on earth or elsewhere as part of routine biological research.

## Methods

### Zag material

Meteorite samples used in this study were obtained from Mr. Rob Wesel, a recognized dealer documented in the *Meteoritical Bulletin* as having handled material that has been formally classified and described in peer-reviewed meteoritical literature^21^.

### Processing and sequencing

The meteorite was UV-irradiated (6 J/cm^2^ at 254 nm) 40 min on each side before taking it into the cleanroom laboratory. In the cleanroom laboratory, the external surface was carefully and mechanically cleaned using a Dremel tool. The sample was wiped with 0.5% bleach solution and followed by Milli-Q® water, using sterile cotton swabs, and then UV-irradiated again, 30 min on each side. A hammer and chisel were used to chip off small pieces from the UV & bleach-treated meteorite sample, which were then pounded to obtain smaller fragments and powder. A Dremel tool with a diamond cutting disc was used to saw off a piece to expose the internal surface of the meteorite sample. A diamond drill bit was then used to shave off powder from the internal surface, which was collected. Two methods of DNA extraction were used on the samples: the Yang-urea extraction method is a modified silica-based method^57^, where 1 M urea was used in place of sodium dodecyl sulfate (SDS) in the extraction buffer. Extraction blanks were included as the negative controls and carried along throughout the whole process. Briefly, the sub-sample was digested with 1 mL of extraction buffer and incubated with rotation at 37 °C for 24 h. Supernatant was collected from the digested sub-sample and concentrated using an Amicon Ultra-4 filter unit (Millipore). The DNA extract was purified with MinElute PCR purification kit (QIAGEN) according to the manufacturer’s instructions and eluted in 110 µL EB buffer. The PowerSoil Pro kit (QIAGEN) was also used, according to the manufacturer’s instructions and eluted in 100 µL Solution C6.

Double-stranded blunt-end libraries for next-generation sequencing were prepared from 20 µL of DNA extracts according to Meyer and Kircher’s protocol^58^ with some modifications. MinElute PCR purification kit (QIAGEN) was used to clean up reactions instead of SPRI beads. The extraction blanks and library blanks using water were included as negative controls and carried along throughout the whole process till the quality control step. The library was quantified by real-time PCR (qPCR) in a 25-µL reaction using Maxima SYBR green master mix (Thermo Fisher Scientific), 200 nM of primer IS7 and 200 nM of primer IS8, to determine the amount of indexing cycles, i.e. Cq-value plus 2 cycles. Indexing PCR amplification was performed in a 50-µL reaction using 6 µL of DNA library, 5 units of AmpliTaq Gold DNA polymerase (Thermo Fisher Scientific), 1߰ GeneAmp Gold buffer (Thermo Fisher Scientific), 2.5 mM MgCl_2_, 250 µM of each dNTP, 200 nM of P7-indexing primer and 200 nM of P5-indexing primer, in duplicates. The PCR reaction was performed at 94 °C for 10 min, 24-28 cycles of (94 °C for 30 s, 60 °C for 30 s, 72 °C for 45 s), and 72 °C for 10 min. The duplicates of the DNA library were pooled and purified with AMPure XP beads (Beckman Coulter). The quality of the library was analyzed by Agilent 2200 Tapestation and the Qubit dsDNA HS assay kit (Invitrogen) was used for DNA quantitation.

DNA libraries from the samples were sequenced along with libraries prepared from no-sample controls from the ancient DNA lab (NSC-aDNA) and pre-PCR lab (NSC-prePCR) to determine the level of background DNA. In total, twenty libraries were prepared and sequenced on one NovaSeq SP lane, 2×150 bp. Sequencing of libraries was performed by SciLifeLab SNP & SEQ Technology Platform. Data analysis was performed by the SciLifeLab Ancient DNA facility.

The lab in which this segment of the work took place follows strict cleaning routines, maintenance, and lab work based on ancient DNA guidelines^59,60^. Personnel wear a disposable body suit with hood, a hair net, a mask with visor, shoe covers, and at least 3 layers of gloves at all times. Entry into the clean laboratories is prohibited if personnel have been in a post-PCR laboratory, unless they have showered and changed into a clean set of clothes. The clean laboratories and equipment are frequently decontaminated with bleach, and/or RNase and DNA AWAY™ Surface Decontaminant (Thermo Scientific), and/or UV irradiation.

Sequencing reads were subjected to adapter trimming, quality filtering, and removal of human-derived sequences. De novo deduplication was performed prior to downstream analyses. To conservatively define sequences associated with internal meteorite material, all reads detected in surface samples (before and after decontamination), extraction blanks, library blanks, and no-sample controls were removed. This subtraction was performed independently for standard-and high-amplification libraries. As an additional conservative step, all sequences present across all material classes were excluded.

### Sequencing data retrieval

The NT database from BLAST was downloaded from NCBI’s FTP site^61^ in FASTA format in December 2023. Subsamples of decreasing fractions (0.1, 0.01, 0.001, 0.0001, 0.00001 and 0.000001) were extracted using SeqKit tool (v2.5.1)^62^. The assembly of a newly discovered microbe was kindly provided by Wild Biotech^27^. From this assembly, 1,000 sequences of length 176 bases were extracted for subsequent analysis. Additionally, random and shuffled sequences were generated with a Python script to produce 1,000 sequences of length 176 bases. The random sequences were generated with a probability of 0.25 for each nucleotide, whereas the shuffled sequences maintained the same nucleotide probability distribution as the “distant-nt” sequences.

### Control datasets

Coding control datasets consisted of sequence fragments sampled in-frame from annotated protein-coding regions of *E. coli* K-12 MG1655. A noncoding control dataset was generated by sampling fragments from intergenic regions of the *E. coli* genome, excluding any overlap with annotated coding sequences. To account for sequencing-related artifacts, an additional *E. coli* control dataset was generated from Illumina reads that failed standard quality-control filters and did not map to the reference genome. These reads were restricted to canonical nucleotides and were used to represent noncoding, error-prone sequence material. Two additional Zag-derived control datasets were constructed from raw Illumina reads prior to nucleotide alignment filtering. One dataset consisted of randomly sampled raw reads, while a second excluded any sequences matching the final internal Zag dataset (or their reverse complements). If necessary, a relaxed exclusion criterion based on shared 31-mers was applied prior to sampling. These datasets were used to test whether observed signals were driven by properties of the raw sequencing pool rather than by the filtered internal dataset. For codon-usage analyses only, additional coding control datasets were generated from annotated coding sequences of *Homo sapiens*, *Saccharomyces cerevisiae*, and *Arabidopsis thaliana*. Fixed-length fragments of 150 nt composed of canonical nucleotides were sampled from RefSeq coding regions. These datasets were analyzed alongside the *E. coli* coding and Zag datasets to assess cross-kingdom codon usage patterns.

### Coding structure and periodicity analysis

Open reading frame structure was assessed by translating sequences in all six frames and quantifying stop codon distributions, including distances between stops and the minimum number of in-frame stops per sequence. Statistical significance was evaluated relative to random-base null models. To assess protein-coding organization, periodicity analyses were performed across all six reading frames. Period-3 signals were evaluated using Z-score based statistics derived from dinucleotide-preserving shuffled controls, as well as using first- and second-order Markov models conditioned on codon phase. Additional analyses were conducted to detect non-canonical periodic structure, including period-4 tests based on phase-specific 2-mer distributions and Jensen–Shannon divergence. Codon usage was analyzed using both 61-codon (sense-only) and 64-codon frameworks. For each sequence, the frame exhibiting the strongest period-3 signal was selected, and codon frequencies were computed. Comparisons across datasets were performed using distance metrics including cosine similarity and Kullback–Leibler divergence. First- and second-order Markov models were trained on pooled coding and noncoding training sets. Log-likelihoods under each model were computed for all sequences, and likelihood ratios were used as classification scores. Performance was assessed using receiver operating characteristic analyses.

### Sequence complexity and motif analysis

Shannon entropy of 3-mers was computed for all sequences based on the equation:

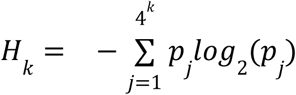

(*k*, *k-*mer length) and compared to dinucleotide-preserving shuffled controls. Homopolymer run lengths were quantified and converted to Z-scores using shuffled null distributions. Inverted-repeat structures were identified by detecting complementary arms within each sequence, with significance assessed relative to shuffled controls. Global enrichment and depletion of 3-mers and 4-mers were evaluated using Z-scores and fold-change statistics.

### Hexamer based structural analysis

Hexamer frequencies were computed for coding and noncoding training datasets and normalized to probability distributions. For each sequence, a hexamer score was calculated as the length-normalized sum of LLR between coding and noncoding hexamer probabilities across all overlapping hexamers.

### Gaussian mixture models

We used Gaussian mixture models (GMMs) to assess the mixture complexity of per-sequence scores in the two-dimensional space of hexamer coding score and 3-periodic Markov log-likelihood ratio (LLR). For each dataset, GMMs with *k* = 1 to 8 components were fitted using full covariance matrices, with five random initializations per *k* (random seed 42). Model selection was performed via the Bayesian Information Criterion (BIC); the number of components *k*\* minimizing BIC was chosen. This procedure was applied separately to each dataset.

### Hartigan’s dip test

Hartigan’s dip test for unimodality was applied to the first principal component (PC1) of the hexamer–Markov score space within each dataset. For each dataset, the two-dimensional feature matrix (hexamer score × Markov LLR) was standardized (zero mean, unit variance), then PCA was performed to obtain two principal components. PC1 was extracted and the dip statistic was computed using the diptest Python package. A bootstrap *p*-value was obtained with 2,000 bootstrap samples (seed 42) to test the null hypothesis of unimodality. Low *p*-values indicated departure from unimodality (e.g. bimodality or multimodality).

### *k*-mer counting

All FASTA files were processed with Jellyfish, a tool designed for counting *k*-mers in a sliding window manner. Jellyfish generates output in a binary format, which can be translated into a human-readable format, queried for specific *k*-mers, and used to produce statistics of the counted *k*-mers. The NT FASTA file was used as input to count *k*-mers of lengths 12, 14, 16, 18, and 20. The representation of *k*-mers was calculated as the number of distinct *k*-mers divided by the number of possible *k*-mers of length *k* (4*^k^*).

### Assembly of foreign sequences

The sequences were constructed using a Python script in an iterative manner, starting with a sample of unrepresented 20-mers from the NT. The initial sample was obtained by extending unrepresented 17-mers from a previous search (results not reported) to 20-mers. This step produced a pool of approximately 38 million unrepresented 20-mers as a starting point. The building algorithm attempts to elongate each *k*-mer in the pool by appending one nucleotide at a time. It then checks whether the newly generated *k*-mer at the 3’ end exists in the pool of unrepresented *k*-mers. If the *k*-mer exists, the algorithm continues with this new *k*-mer and repeats the one-base elongation process until the longest possible sequence is reached. If the *k*-mer does not exist, the elongation stops and reinitiates with the next *k*-mer.

*k*-mers that are missing from the unrepresented pool are either present in the NT database or absent from the current sample pool. Therefore, when the elongation process stops at a length close to the current longest sequence, the missing *k*-mers are queried in the Jellyfish 20-mer count file. The resulting *k*-mers that are truly missing from the NT database are added to the sample pool, and another search iteration begins.

### Phylogenetic analysis

The phylogenetic tree of life was obtained from Nature Microbiology (reference: https://www.nature.com/articles/nmicrobiol201648). Records assigned to each genus on the tree were retrieved from a 10% subsample of the NT FASTA file using the SeqKit tool (v2.5.1), and their *k*-mers were counted using the Jellyfish tool. The visualization of the tree and the log representation of *k*-mers was performed using iTOL (v6.9.1)^63^.

### Technical artifact tests

Chimera detection: reads were restricted to the most common length (mode length) to form a same-length cohort. The top 50 most abundant unique sequences (by exact count) were defined as “parents.” Each read was split at the midpoint into left and right halves. A read was classified as a chimera if (i) the best-matching parent for the left half (by Hamming distance, d ≤ 2) differed from the best-matching parent for the right half, and (ii) both halves were within Hamming distance 2 of their respective best parents. The observed chimera fraction was compared to a null distribution generated by permuting right halves among reads (30 permutations). The Z-score of the observed chimera fraction vs. the null mean and standard deviation was reported.

Error halo: parent sequences were defined as unique sequences with count ≥ 2. Child sequences were those within Hamming distance ≤ 3 of their nearest parent (from the top 200 parents by abundance). The fraction of reads in the “error halo” (reads in sequences assigned to a parent at d ≤ 3) was computed. Counts at distances 0, 1, 2, and 3 from the assigned parent were also recorded. This metric captures the extent to which reads cluster around abundant “parent” sequences, as expected under PCR amplification with a low per-base error rate.

Rank abundance and duplicate metrics: exact duplicate counts were used to compute: (i) the fraction of reads in the single most abundant sequence (top-1 fraction), (ii) the Gini coefficient of the abundance distribution, (iii) Shannon entropy of the abundance distribution, and (iv) Hill number of order 1 (exp(entropy)). These metrics summarize the “jackpot” or amplification-skew character of the dataset.

Family clustering: sequences of length ≥ 60 nt were used. A 40-nt seed (positions 20–60) was extracted from each sequence. Sequences were clustered by greedy assignment: each sequence was assigned to the first existing centroid within Hamming distance ≤ 3 of its seed, or to a new centroid if none matched. Family size statistics (mean, median, 95th percentile, max) and the fraction of reads in families of size ≥ 5 were reported.

Per-cycle drift: for each cycle (position), the base composition (A, C, G, T fractions) was computed across reads. The mean L1 deviation of per-position composition from the global composition (over positions present in ≥ 80% of reads) was computed. Linear slopes of G and C fractions versus position were also computed to capture cycle-dependent bias.

Adapter/primer matching: To identify demonstrable technical sequence carryover without assuming biological sequence structure, reads were screened exclusively against known Illumina, TruSeq, and Nextera adapter and primer sequences using a two-stage procedure. First, a *k*-mer–based screen was performed with BBDuk (BBTools) against a curated reference set of 167 adapter/primer sequences using a fixed *k*-mer length of 23 nucleotides, allowing up to one mismatch and considering both forward and reverse-complement orientations. No low-complexity, homopolymer, or composition-based filtering was applied.

| Method | Purpose | Implementation |
| --- | --- | --- |
| GMM + BIC | Mixture complexity of hexamer × Markov scores | sklearn GaussianMixture, K = 1–8, BIC selection |
| Hartigan’s dip test | Unimodality of PC1 | dip test package, 2000 bootstrap |
| PCR chimera | Split chimeras (left/right parent mismatch) | Top-50 parents, $d \leq 2$, 30 null permutations |
| Error halo | Reads within $d \leq 3$ of parents | Top-200 parents, Hamming assignment |
| Rank-abundance | Jackpot / duplicate distribution | Gini, Shannon entropy, top-1 fraction |
| Per-cycle drift | Base composition vs position | L1 deviation, G/C slope |
| Adapter/primer screening | Technical oligo match | Illumina + Nextera refs |

### MinHash Jaccard similarity

To quantify similarity between sequence datasets while minimizing sensitivity to sequence length and abundance, MinHash-based Jaccard similarity was computed on *k*-mer presence–absence profiles. For each sequence, the set of distinct *k*-mers (*k* = 21) was enumerated, and MinHash sketches of fixed size were generated using a consistent random seed. Pairwise Jaccard similarities were then estimated between the datasets. To establish null expectations, dinucleotide-preserving shuffled versions of each dataset were generated and analyzed in parallel. Estimated Jaccard similarities were summarized across replicate sketches, and comparisons between observed and shuffled datasets were used to assess whether the Zag-derived sequences exhibited non-random overlap with known biological sequence space.

## Supporting information

Media guidance

## Acknowledgements

Processing of DNA and data analysis were performed by the SciLifeLab Ancient DNA facility. Sequencing was performed by the SNP & SEQ Technology Platform in Uppsala, part of the National Genomics Infrastructure (NGI) Sweden and Science for Life Laboratory. The SNP & SEQ Platform is also supported by the Swedish Research Council and the Knut and Alice Wallenberg Foundation. The authors wish to thank Joe Davis for valuable ideas and discussions; Dr. Haider Hassan and the team at Bridge Informatics for their technical expertise, valuable discussions, and work carried out in this project; Dr. David Tolpin for valuable help with Bayesian analysis of the data; Mr. Itay Rusinek for critical reading of the manuscript and tests; and Mr. Marius Nacht for his generosity and vision.

## Author contributions

C. F., G. M. C., and I. B. designed the study, analyzed data, and wrote the manuscript.

## Declaration of competing interests

The authors declare no competing interests.

---

**Table S1.** Test summary.

| # | Test | Finding | Figure |
| --- | --- | --- | --- |
| 1 | Base composition | A 28%, T 29%, G 23%, C 20%. |  |
| 2 | Translation in all 6 frames | Gaps between stop codons follow a near-random decay pattern; no widespread long coding regions |  |
| 3 | Stop codon avoidance | No evidence of systematic coding-like stop depletion across the dataset | 2C |
| 4 | 3-mer Shannon entropy vs mono- and dinucleotide preserving shuffles | Lower entropy than expected in a random sequence | 3A, 3B, 3C |
| 5 | Triplet/quadruplet periodicity | No periodicity found in all 6 frames | 2A, 2E |
| 6 | Frame-aware hexamer coding score | No coding structure | 2B |
| 7 | Periodicity scan (p=2 to p=8) | No periodicity | 2D |
| 8 | A more sensitive quadruplet periodic structures | None found | 2F |
| 9 | 3- and 4-mer enrichment/depletion | Enrichment in simple homopolymers and tandem fragments | 3D, 3E |
| 10 | Palindrome/hairpin enrichment | No systematic enrichment found | 3F |
| 11 | Markov 1st and 2nd order LLR | No markov grammar found (LLR = negative) | 4A, 4B |
| 12 | GMM component count by BIC | Dataset is unimodal | 5A |
| 13 | Hartigan's dip test on PC1 | Dataset is unimodal | 5B |
| 14 | Hexamer vs Markov space clustering | Dataset contains multiple manifolds consistent with heterogenous, non-grammatical sequence generation | 4C, 4D, 4E |
| 15 | PCR chimeras | <0.1% |  |
| 16 | PCR error halo | Most reads have distance of 0; no sharp spike at distance 1; distances 1-3 not dominated by a few parent sequences |  |
| 17 | PCR error halo scatter | No amplification-driven family expansion; parents have maximum 1-2 exact matches |  |
| 18 | PCR rank abundance | Rank 1-2: a count of 2. All others are singletons. No power-law or geometric decay |  |
| 19 | PCR jackpot | Top 1 read fraction = 0.031%; top 10 fraction = 0.19%; top 100 fraction = 1.59%; Gini coefficient = 0.014 |  |
| 20 | Matching sequences traceable to a reference set of technical oligonucleotides | Exists in ~ 13% sequences out of 17,000 ($k=23$); upper bound of ~ 33% ($\min k=10$). | S1 |
| 21 | MinHash Jaccard | Jaccard similarity ~ 0.0007 |  |
| 22 | Mean Phred Q (per read) | 17k matched reads: mean Q = 36.32, median = 36.67 (min 28.5, max 37.0); high quality, not consistent with low-quality or adapter-dimer artifact. |  |
| 23 | Per-base quality profile | Mean Q at pos 1/75/151 = 36.4, 36.4, 35.3 |  |
| 24 | Contamination (FastQ Screen, lane-level) | Sample lanes: No hits 39.20%, Human 36.95%, Adapters 45.01%; Extraction blank: 38.91%, 34.16%, 43.97%. Similar; no evidence 17k lanes are more contaminated than Extraction blank. |  |

**Figure S1.**
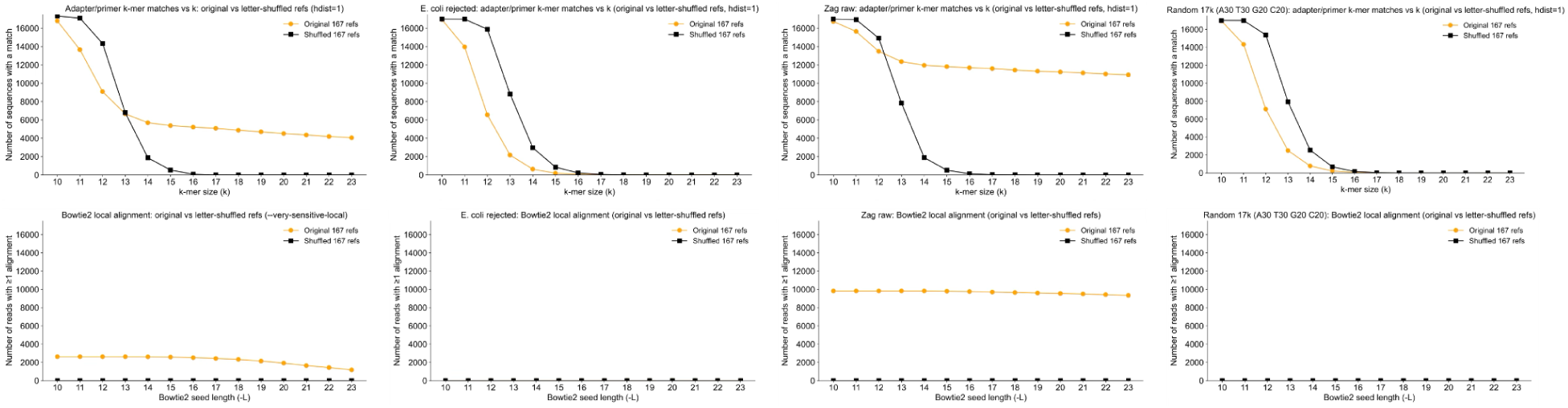
Adapter/primer *k*-mer match sweep: original versus letter-shuffled reference controls. Number of reads with at least one k-mer match (*k*=23 down to 10, hdist=1) to 167 Illumina/Nextera adapter/primer sequences (orange) versus letter-shuffled references preserving composition (black). At *k*=23: 4,052 reads (23.4%) matched original refs; 0 matched shuffled. At *k*≥15, shuffled control remains near zero; plateau in original curve (*k*=15–23) reflects genuine adapter/primer signal. Surge at *k*<15 reflects loss of specificity (chance matches). Real contamination estimated at ∼5,000–5,500 (*k*=15–16 cutoff). Bowtie2 high-confidence confirmation (≥20 bp, ≥98% identity, end-anchored): 169 reads (0.97%) disqualified; 17,166 (99.03%) retained. Upper bound under permissive (mink=10) screen ∼33%. MinHash Jaccard similarity (main dataset vs dinucleotide-shuffled) ∼0.0007, several orders of magnitude below null expectation.

## Notes

### Competing Interest Statement

The authors have declared no competing interest.

### Summary of Updates

This version has been revised to report >20 analytical and statistical tests performed on the source sequences, placing the possibility of technical artifacts or metagenomic contamination at a low likelihood.

## References

1. Wiens, J. J. How many species are there on Earth? Progress and problems. PLoS Biol. 21, e3002388 (2023).

2. Camacho, C. et al. BLAST+: architecture and applications. BMC Bioinformatics 10, 421 (2009).

3. Wood, D. E., Lu, J. & Langmead, B. Improved metagenomic analysis with Kraken 2. Genome Biol 20, 257 (2019).

4. Ledvina, H. E. & Whiteley, A. T. Conservation and similarity of bacterial and eukaryotic innate immunity. Nat. Rev. Microbiol. 22, 420–434 (2024).

5. Takemata, N., Samson, R. Y. & Bell, S. D. Physical and Functional Compartmentalization of Archaeal Chromosomes. Cell 179, 165–179.e18 (2019).

6. Alvarez-Ponce, D. & McInerney, J. O. The human genome retains relics of its prokaryotic ancestry: human genes of archaebacterial and eubacterial origin exhibit remarkable differences. Genome Biol. Evol. 3, 782–790 (2011).

7. Isenbarger, T. A. et al. The most conserved genome segments for life detection on Earth and other planets. Orig. Life Evol. Biosph. 38, 517–533 (2008).

8. Moody, E. R. R. et al. The nature of the last universal common ancestor and its impact on the early Earth system. Nat Ecol Evol (2024) doi:10.1038/s41559-024-02461-1.

9. Ban, N. et al. A new system for naming ribosomal proteins. Curr. Opin. Struct. Biol. 24, 165–169 (2014).

10. Grossman, J. n. The Meteoritical Bulletin, No. 83, 1999 July. Meteorit. Planet. Sci. 34, A169–A186 (1999).

11. Gaffey, M. J. & Gilbert, S. L. Asteroid 6 Hebe: The probable parent body of the H-type ordinary chondrites and the IIE iron meteorites. Meteorit. Planet. Sci. 33, 1281–1295 (1998).

12. [No title]. https://arxiv.org/pdf/1705.10515.

13. [No title]. https://www.lpi.usra.edu/meetings/metsoc99/pdf/5050.pdf.

14. Whitby, J., Burgess, R., Turner, G., Gilmour, J. & Bridges, J. Extinct (129)I in halite from a primitive meteorite: evidence for evaporite formation in the early solar system. Science 288, 1819–1821 (2000).

15. Gilmour, J. D., Pravdivtseva, O. V., Busfield, A. & Hohenberg, C. M. The I-Xe chronometer and the early solar system. Meteorit. Planet. Sci. 41, 19–31 (2006).

16. Chan, Q. H. S. et al. Organic matter in extraterrestrial water-bearing salt crystals. Sci Adv 4, eaao3521 (2018).

17. Kebukawa, Y. et al. A novel organic-rich meteoritic clast from the outer solar system. Sci. Rep. 9, 3169 (2019).

18. Complex mixture of organic matter in a xenolithic clast from the Zag meteorite revealed by coordinated analyses using AFM-IR, NanoSIMS and STXM/XANES. Icarus 400, 115582 (2023).

19. Kvenvolden, K. et al. Evidence for extraterrestrial amino-acids and hydrocarbons in the Murchison meteorite. Nature 228, 923–926 (1970).

20. Meierhenrich, U. J., Muñoz Caro, G. M., Bredehöft, J. H., Jessberger, E. K. & Thiemann, W. H.-P. Identification of diamino acids in the Murchison meteorite. Proc. Natl. Acad. Sci. U. S. A. 101, 9182–9186 (2004).

21. Callahan, M. P. et al. Carbonaceous meteorites contain a wide range of extraterrestrial nucleobases. Proc. Natl. Acad. Sci. U. S. A. 108, 13995–13998 (2011).

22. Extraterrestrial nucleobases in the Murchison meteorite. Earth Planet. Sci. Lett. 270, 130–136 (2008).

23. Oba, Y. et al. Identifying the wide diversity of extraterrestrial purine and pyrimidine nucleobases in carbonaceous meteorites. Nat. Commun. 13, 2008 (2022).

24. Dabney, J., Meyer, M. & Pääbo, S. Ancient DNA damage. Cold Spring Harb Perspect Biol 5, (2013).

25. Sayers, E. W. et al. Database resources of the national center for biotechnology information. Nucleic Acids Res. 50, D20–D26 (2022).

26. Bussi, Y., Kapon, R. & Reich, Z. Large-scale k-mer-based analysis of the informational properties of genomes, comparative genomics and taxonomy. PLoS One 16, e0258693 (2021).

27. Levin, D. et al. Diversity and functional landscapes in the microbiota of animals in the wild. Science 372, (2021).

28. Monzoorul Haque, M., Ghosh, T. S., Komanduri, D. & Mande, S. S. SOrt-ITEMS: Sequence orthology based approach for improved taxonomic estimation of metagenomic sequences. Bioinformatics 25, 1722–1730 (2009).

29. Ghosh, T. S., Monzoorul Haque, M. & Mande, S. S. DiScRIBinATE: a rapid method for accurate taxonomic classification of metagenomic sequences. BMC Bioinformatics 11 Suppl 7, S14 (2010).

30. Gerlach, W. & Stoye, J. Taxonomic classification of metagenomic shotgun sequences with CARMA3. Nucleic Acids Res 39, e91 (2011).

31. A survey of k-mer methods and applications in bioinformatics. Comput. Struct. Biotechnol. J. 23, 2289–2303 (2024).

32. Ounit, R., Wanamaker, S., Close, T. J. & Lonardi, S. CLARK: fast and accurate classification of metagenomic and genomic sequences using discriminative k-mers. BMC Genomics 16, 236 (2015).

33. Ondov, B. D. et al. Mash: fast genome and metagenome distance estimation using MinHash. Genome Biol. 17, 132 (2016).

34. Lee, R. C. & Ambros, V. An extensive class of small RNAs in Caenorhabditis elegans. Science 294, 862–864 (2001).

35. Sim, G., et al. Determining the defining lengths between mature microRNAs/small interfering RNAs and tinyRNAs. bioRxiv (2023) doi:10.1101/2023.10.27.564437.

36. Li, Z. et al. Characterization of viral and human RNAs smaller than canonical MicroRNAs. J. Virol. 83, 12751–12758 (2009).

37. Li, Z. et al. Extensive terminal and asymmetric processing of small RNAs from rRNAs, snoRNAs, snRNAs, and tRNAs. Nucleic Acids Res. 40, 6787–6799 (2012).

38. Baldrich, P. et al. Plant Extracellular Vesicles Contain Diverse Small RNA Species and Are Enriched in 10- to 17-Nucleotide ‘Tiny’ RNAs. Plant Cell 31, 315–324 (2019).

39. Sim, G. et al. Manganese-dependent microRNA trimming by 3’→5’ exonucleases generates 14-nucleotide or shorter tiny RNAs. Proc. Natl. Acad. Sci. U. S. A. 119, e2214335119 (2022).

40. Turk, R. M., Chumachenko, N. V. & Yarus, M. Multiple translational products from a five-nucleotide ribozyme. Proc. Natl. Acad. Sci. U. S. A. 107, 4585–4589 (2010).

41. Yarus, M. The meaning of a minuscule ribozyme. Philos. Trans. R. Soc. Lond. B Biol. Sci. 366, 2902–2909 (2011).

42. Kim, S. C., O’Flaherty, D. K., Giurgiu, C., Zhou, L. & Szostak, J. W. The Emergence of RNA from the Heterogeneous Products of Prebiotic Nucleotide Synthesis. J Am Chem Soc 143, 3267–3279 (2021).

43. Xu, J. et al. Selective prebiotic formation of RNA pyrimidine and DNA purine nucleosides. Nature 582, 60–66 (2020).

44. Roberts, S. J. et al. Selective prebiotic conversion of pyrimidine and purine anhydronucleosides into Watson-Crick base-pairing arabino-furanosyl nucleosides in water. Nat Commun 9, 4073 (2018).

45. Xu, J., Green, N. J., Russell, D. A., Liu, Z. & Sutherland, J. D. Prebiotic Photochemical Coproduction of Purine Ribo- and Deoxyribonucleosides. J Am Chem Soc 143, 14482–14486 (2021).

46. Neubeck, A. & McMahon, S. Prebiotic Chemistry and the Origin of Life. (Springer Nature, 2022).

47. Smoukov, S. K., Seckbach, J. & Gordon, R. Conflicting Models for the Origin of Life. (John Wiley & Sons, 2023).

48. Lambert, J. B., Gurusamy-Thangavelu, S. A. & Ma, K. The silicate-mediated formose reaction: bottom-up synthesis of sugar silicates. Science 327, 984–986 (2010).

49. Furukawa, Y., Horiuchi, M. & Kakegawa, T. Selective stabilization of ribose by borate. Orig. Life Evol. Biosph. 43, 353–361 (2013).

50. Ricardo, A., Carrigan, M. A., Olcott, A. N. & Benner, S. A. Borate minerals stabilize ribose. Science 303, 196 (2004).

51. Hirakawa, Y., Kakegawa, T. & Furukawa, Y. Borate-guided ribose phosphorylation for prebiotic nucleotide synthesis. Sci. Rep. 12, 1–7 (2022).

52. Stephenson, J. D., Hallis, L. J., Nagashima, K. & Freeland, S. J. Boron enrichment in martian clay. PLoS One 8, e64624 (2013).

53. Mason, B. H. Handbook of Elemental Abundances in Meteorites. (Gordon & Breach Publishing Group, 1971).

54. Boron in chondritic meteorites. Geochim. Cosmochim. Acta 52, 2311–2319 (1988).

55. Krishnamurthy, R., Goldman, A. D., Liberles, D. A., Rogers, K. L. & Tor, Y. Nucleobases in Meteorites to Nucleobases in RNA and DNA? J Mol Evol 90, 328–331 (2022).

56. Bhadra, S., Maranhao, A. C., Paik, I. & Ellington, A. D. One-Enzyme Reverse Transcription qPCR Using Taq DNA Polymerase. Biochemistry 59, 4638–4645 (2020).

57. Yang, D. Y., Eng, B., Waye, J. S., Dudar, J. C. & Saunders, S. R. Technical note: improved DNA extraction from ancient bones using silica-based spin columns. Am J Phys Anthropol 105, 539–543 (1998).

58. Meyer, M. & Kircher, M. Illumina sequencing library preparation for highly multiplexed target capture and sequencing. Cold Spring Harb Protoc 2010, db.prot5448 (2010).

59. Fulton, T. L. & Shapiro, B. Setting Up an Ancient DNA Laboratory. Methods Mol Biol 1963, 1–13 (2019).

60. Shapiro, B. A. & Hofreiter, M. Ancient DNA: Methods and Protocols. (2012).

61. Index of /blast/db/FASTA. https://ftp.ncbi.nlm.nih.gov/blast/db/FASTA.

62. Shen, W., Le, S., Li, Y. & Hu, F. SeqKit: A Cross-Platform and Ultrafast Toolkit for FASTA/Q File Manipulation. PLoS One 11, e0163962 (2016).

63. Website. https://academic.oup.com/bioinformatics/article/23/1/127/188940.

