## Supplementary material for "Composition and higher-order structure in nucleic acids sequenced from a chondrite": Media guidance

### What this study reports

In this study, we report to have successfully read the sequence of a small amount of nucleic acids from inside the Zag meteorite, a rock that fell to Earth in Morocco in 1998. A battery of tests assigned a low likelihood to contamination or technical artifacts as explanations for this material. We then asked whether these sequences resemble the DNA of a living organism. They do not. They carry no detectable structure of a biological language or grammar. They look like polymers that were created chemically from available building blocks under certain constraints — not random, but not a code either.

Nucleic acids such as DNA are regular chemicals. Their building blocks are known to form spontaneously under the right conditions, and several of them have already been identified inside meteorites by other groups. It is now the scientific consensus that nucleic acids first formed on Earth through chemistry alone, before life began. So the presence of nucleic acid material in a meteorite, while noteworthy, is not by itself evidence of life.

We do not claim to have found the DNA of an extraterrestrial organism. We report the sequencing and characterization of nucleic acid material recovered from a meteorite, and we encourage independent replication of our findings.

### Errors to avoid

We and our scientific peers would regard each of the following as factual errors:

- **"Alien DNA" / "Extraterrestrial life"** — We found nucleic acid chemistry, not an organism. The sequences fail every test for biological signatures.
- **"Scientists decoded a genome"** — There is no genome. These are short sequence fragments with no coding organization.
- **"DNA from the outer solar system"** — The Zag meteorite originates from the inner asteroid belt.
- **"They proved it is not contamination"** — We did not prove this. We estimate a low likelihood of contamination.
- **"Confirmed" / "Landmark discovery"** — This is a single study. Confirmation requires independent replication by other groups.

For queries: Ido Bachelet
